# Unbalanced dietary patterns contribute to the pathogenesis of precocious puberty by affecting gut microbiota and host metabolites

**DOI:** 10.1101/2021.04.07.438759

**Authors:** Ying Wang, Dingfeng Wu, Hongying Li, Xiangrong Liang, Na Jiao, Wenxing Gao, Lu Zhao, Han Yu, Qian Wang, Yongsheng Ge, Changying Zhao, Meiling Huo, Ruifang Cao, Sheng Gao, Liwen Tao, Yunchao Ling, Lingna Zhao, Xin Lv, Yi Liu, Lehai Zhang, Haokui Zhou, Guoqing Zhang, Guoping Zhao, Lei Zhang, Ruixin Zhu, Zhongtao Gai

## Abstract

Precocious puberty (PP) mostly stems from endocrine disorders. However, its triggering factors, especially for the early onset of partial PP, and the associated pathogenic mechanisms remain ambiguous. In this study, a systematic analysis in the form of a questionnaire of lifestyles, gut microbiome, and serum metabolome data was carried out to examine the pathogenesis of PP in a cohort comprised of 200 girls, with or without PP. The analysis revealed substantial alterations in gut microbiota, serum metabolites, as well as lifestyle patterns in the PP group, which were characterized by an elevated abundance of β-glucuronidase-producing and butyrate-producing bacteria, and excessive lipid concentration with decreased levels of organic nitrogen compounds in the serum of the participants. These differential microbes and metabolites tend to be reliable non-invasive diagnostic biomarkers aiding the early diagnosis of PP and exhibit a strong discriminative power (AUC = 0.93 and AUC = 0.97, respectively). Furthermore, the microbial biomarkers were confirmed in an independent validation cohort (n = 83, AUC = 0.85). Moreover, structural equation modeling revealed that unhealthy dietary habits were the primary contributors for the alteration of gut microbiota and serum metabolites, triggering the imbalance in the host hormones that leads to premature physical development. Our study determines a causal relationship among the gut microbiota, host metabolites, diet, and clinical characteristics of preadolescent girls who experienced early onset of PP, and formulates non-invasive diagnostic tools demonstrating excellent performance for the early detection of PP.

## INTRODUCTION

Precocious puberty (PP) refers to the premature occurrence of pubertal development or secondary sexual characteristics, before the age of 8 in girls and 9 in boys, depicted by features such as advanced breast and ovary development along with rapid bone growth or maturation (Root, 2000), where the morbidity rate is also seen to rise progressively (Kim et al., 2015). PP is a hormonal condition predominantly seen in females and can be attributed to endocrine disorders, accompanied by an elevated sex hormone secretion (Du et al., 2019). However, there still exists lack of clarity about the triggering factors of this condition, especially for the premature onset of partial PP, and the pathogenic mechanisms associated with it. It has been reported that dietary patterns seem to considerably influence the estrogen metabolism mechanism, which is inextricably linked with PP (Chen et al., 2018; Kim et al., 2011b; Merzenich et al., 1993; Rogers et al., 2010). Over-nutrition or hyperalimentation, the excessive consumption of processed and high-fat diet, is considered to be the principal agent responsible for the secular decline in pubertal age (Muir, 2006; Soliman et al., 2014). Certain animal studies have suggested that postnatal over-nutrition tends to invariably escalate the secretion of luteinizing hormone (LH), follicle stimulating hormone (FSH), leptin, and insulin levels in pubertal females, while the consumption of postnatal high-fat diet after commencing weaning stimulates premature puberty in females (Soliman et al., 2014). At the same time, harmful dietary patterns seem to significantly affect the composition of human gut microbiota and metabolome (Kong et al., 2014; Sheflin et al., 2017). A number of former studies conducted on adults with estrogen-mediated diseases, such as experiencing menopausal symptoms, revealed that gut microbiota is capable of effectively regulating metabolism and transforming estrogen-like compounds to biologically active forms (Baker et al., 2017; Frankenfeld et al., 2014). Hence, PP has been understood to be the outcome of early activation of hypothalamic-pituitary-gonadal (HPG) axis initiated by certain pathophysiological stimuli, such as gut microbiota or diet patterns (Brito et al., 1999; Cussotto et al., 2018; Qi et al., 2012).

Therefore, we investigated the gut microbiome and serum metabolome with respect to the lifestyle information and clinical characteristics for a cohort of 200 participants, where 133 girls experienced partial PP in early onset stage while 67 were healthy girls. Our study registered an imperative association between the key metabolomics and bacterial biomarkers, and their promising discriminatory power values by analyzing the discovery and validation cohorts. Our innovative structural equation modeling (SEM) analysis demonstrated the direct and/or indirect causal relationships across various factors of this study for determining PP in preadolescent girls, such as gut microbiota, host metabolism, lifestyles, and clinical characteristics.

## RESULTS

### Baseline Characteristics of Participants

200 female participants were recruited for this study (Table 1). 168 stool samples from participants were collected for 16S rRNA sequencing to probe the microbiota alterations existing among girls with PP (n = 105) and normal girls (n = 63). The average age of girls in the PP and the normal groups at the time of stool sample collection was 6.641 (95 confidence interval, ci95 = 0.403) and 7.008 (ci95 = 0.515) years (*P* = 0.278, Table 1), respectively. 129 serum samples were collected for untargeted metabolomics analysis, which included 45 PP and 84 normal girls illustrating an average age of 6.662 (ci95 = 0.458) and 6.250 (ci95 = 0.571) years (*P* = 0.289, Table 1), respectively. Additionally, lifestyle information of the 200 participants was obtained by means of a questionnaire involving 117 variables of dietary patterns, living environment, maternal health, childbirth, and the physical condition of participants as well as their parents. For the dietary patterns of the participants, 15 variables, including seafood (FDR = 2.36e-5), freshwater products (FDR = 2.91e-5), tubers (FDR = 0.65e-3), and vegetables (FDR = 0.0021), showed significant differences between PP and the normal group (Table S1). Likewise, 3 variables depicting the physical condition of the participants, like dental care (FDR = 0.021), eczema (FDR = 0.045), and normal vaccination (FDR = 0.047), also showed significant differences among the two groups under consideration (Table S1).

**Table 1.** The number of sample and age distribution in this study.

|  | 16S (n=168) | Metabolism (n=129) | Overlap (n=97) |
| --- | --- | --- | --- |
| Normal (n=67) | 7.008 ± 0.515 (n=63) | 6.250 ± 0.571 (n=45) | 6.211 ± 0.629 (n=41) |
| PP (n=133) | 6.641 ± 0.403 (n=105) | 6.662 ± 0.458 (n=84) | 6.890 ± 0.530 (n=56) |
| *P* (t-test) | 0.278 | 0.289 | 0.108 |

Age Statistics (mean ± 95 confidence interval) and difference analysis (t-test)

### Gut Microbiota Dysbiosis in Girls Suffering from PP

Gut microbial composition displayed a huge variation between PP and the normal group. As compared to the normal girls (*P* < 0.001, Fig. 1A), significantly elevated bacterial richness was observed in the PP group. At the same time, the microbial diversity in the PP group appeared to be substantially distinct from that of the normal group, which was further validated by PERMANOVA test (pseudo-F = 3.24, *P* = 0.001, Fig. 1B). 45 notably differential amplicon sequence variants (ASVs) were identified from the gut microbiota samples of the PP group and the normal controls (FDR < 0.05, Table S2). The abundance of ASVs, assigned as Bacteroidaceae, Ruminococcaceae, Faecalibacterium, Enterobacteriaceae, and Escherichia-Shigella, was witnessed to have increased, while those of genus *Agathobacter* and family Peptostreptococcaceae (*Romboutsia* and *Intestinibacter (1)*) seemed to have decreased in the PP group (Fig. 1D, Table S2). Most of these differential taxa exhibited the potential of encoding/producing β-glucuronidase (Fig. 1D, Table S2), an enzyme that deconjugates estrogens into their active forms (Mcintosh et al.).

**Fig. 1.**
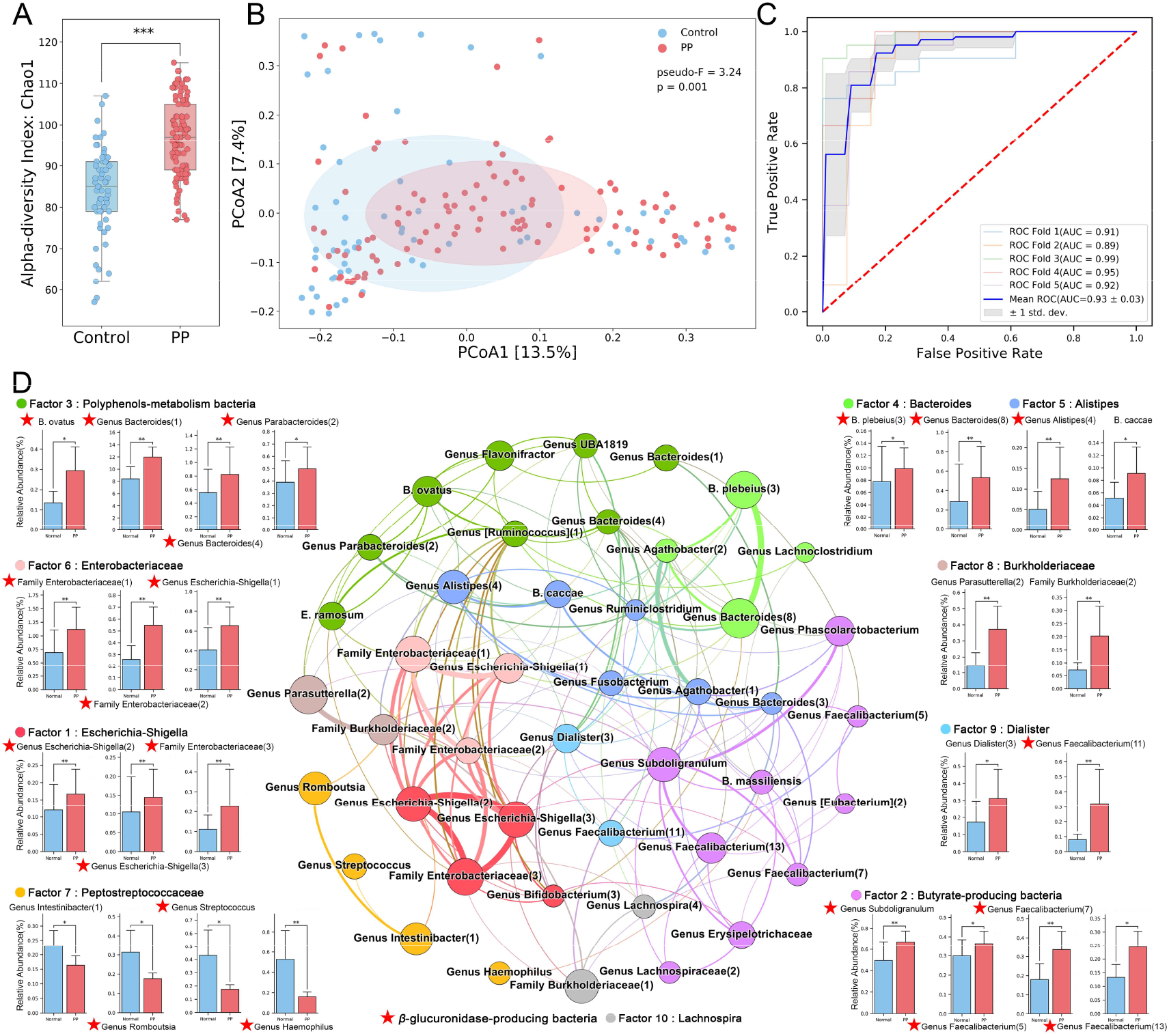
Gut microbiota dysbiosis in girls with partial PP. A. The α diversity of gut microbiota based on Chao1 for PP and normal group (***: *P*< 0.001). B. PCoA of bacterial beta diversity based on the Bray-Curtis distance between PP and normal groups. C. The ROC curve of the disease discriminating ability with 45 differential ASVs. D. Co-occurrence network of differential microbiota through Spearman’s rank correlation analysis with *P*<0.05. Microbiota are colored by their main latent microbial factor and the strength of correlation is represented by line thickness. The abundance changes of the representative microbiota is displayed as bar plot (*: FDR < 0.05, **: FDR < 0.01). β-glucuronidase-producing bacteria are labeled with a red star.

Furthermore, alterations on microbiome-mediated functional potentials were also explored, which led to the identification of 88 differential pathways (FDR < 0.05) between PP and the normal group (Fig. S1A). The PP group demonstrated enhanced activity levels in most metabolic processes, such as metabolizing carbohydrate, cofactor and vitamin, fatty acid and lipid, and inorganic nutrient metabolism.

### Microbial Markers Could Act as Non-Invasive Tools for PP Diagnosis

To explore the possibility of differential microbes functioning as prospects of novel non-invasive tools for PP diagnosis, a classification model was designed by employing 45 differential ASVs via random forest algorithm. The model emerged highly capable for performing the clinical diagnosis of PP, with an area under the receiver operating characteristic curve (AUC) of about 0.93 (Fig. 1C). Afterwards, the classification model was validated in an independent cohort (n = 83, Table S3) to further measure its generalization ability. The model constituting of 13 out of 45 differential ASVs that exhibited the same abundance change pattern accomplished excellent results for distinguishing between PP and the normal group with an AUC of about 0.85 (Fig. S2B). These 13 ASVs primarily belonged to *Bacteroides* and Enterobacteriaceae (Table S3) and illustrated critical elevation regarding several metabolic processes in the PP group (Fig. S1B).

### Latent Microbial Factors Revealed Underlying Relationships of Altered Microbiome

Delving into the underlying relationships among differential taxa, we performed exploratory factor analysis (EFA) and identified 10 latent microbial factors, which revealed the latent patterns of the change in gut microbiota. These factors were then presented in the microbial co-occurrence network (Fig. 1D). The classification models designed on the basis of these 10 microbial factors achieved a high accuracy (AUC = 0.88, Fig. S2A) in distinguishing the PP group from the normal controls, not only indicating the efficacy of the EFA analysis but also preserving the vital information possessed by differential taxa. Methodically speaking, Gammaproteobacteria, such as family Enterobacteriaceae and genus *Escherichia-Shigella*, which retained a strong correlation with each other, were members of Factor 1 and 6 (Fig. 1D). The chief members of Factor 2, genus *Subdoligranulum* and *Faecalibacterium* of family Ruminococcaceae, are known as butyrate-producing bacteria in gut microbiota (Cussotto et al., 2018). Besides, four taxa, *Romboutsia, Intestinibacter, Streptococcus*, and *Haemophilus*, present in a considerably low concentration in the PP group, expressed a strong loading in Factor 7 (Fig. 1D, Table S2). Notably, several species (except Peptostreptocaccaceae, Factor 7), were witnessed in elevated measures in the PP group (Fig. 1D, Table S2). Among them, factor *Bacteroides*, butyrate-producing bacteria, Enterobacteriaceaes, and Burkholderiaceae appeared to be the most relevant towards the microbial metabolic pathways (Fig. S1B) and could prove to be instrumental in microbial biological functions.

### Altered Serum Metabolome in Girls Suffering from PP

Untargeted metabolomic profiles were evaluated under LC-MS/MS system in 84 PP and 45 normal girls (Table 1) for the purpose of investigating metabolite alterations, and 182 differential metabolites were identified (FDR < 0.05, Table S4). Among them, 131 differential metabolites, such as Phenylalanine, 4-Guanidinobutamide, and Lysophosphatidylcholine (LPC), were present in decreased quantities in the PP group as compared to the normal controls (Table S4). Significantly, these 182 differential metabolites were remarkably efficient in differentiating the PP group from the normal controls and obtained an AUC of 0.97 (Fig. 2B), which appears to be higher than microbial markers.

**Fig. 2.**
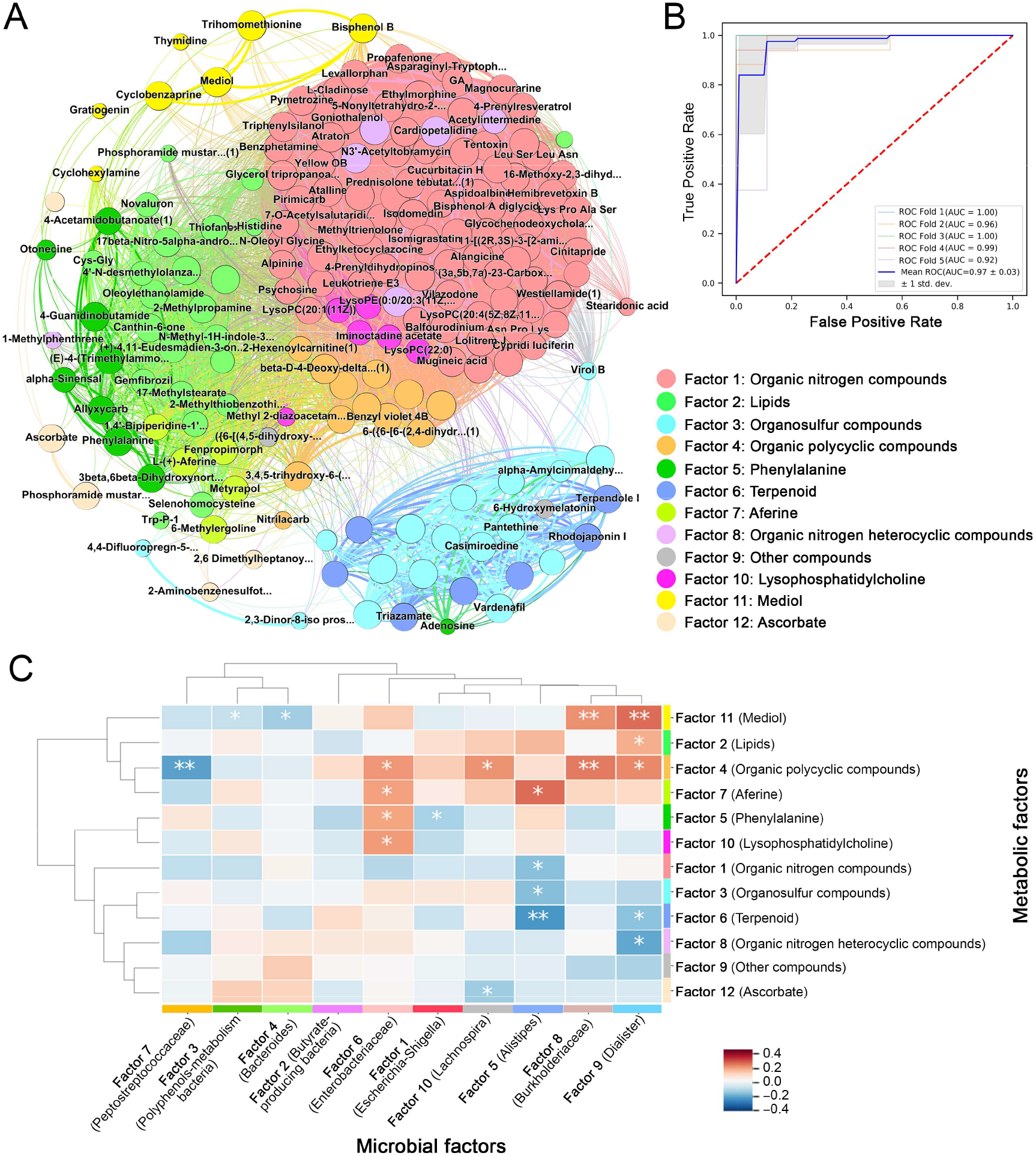
The change of serum metabolome in girls with partial PP. A. Co-occurrence network of differential metabolites through Spearman’s rank correlation analysis with *P*<0.05. Metabolites are colored by their main latent metabolic factor and the strength of correlation is represented by line thickness. B. The ROC curve of the disease discriminating ability with 182 differential metabolites. C. Heatmap plot of Pearson correlation between microbial and metabolic latent factors. (*: *P* < 0.05, **: *P* < 0.01.)

### Organic Nitrogen Compounds and Lipids Were the Characteristic Latent Metabolic Factors

Analogous to the EFA analysis for microbiota, 12 latent metabolic factors were identified based on their differential metabolites (Fig. 2A, Table S4). Preserving the foremost information, the metabolic factors exhibited the detection capability comparable to that of the differential metabolites for detecting PP (AUC = 0.97, Fig. S2C). Specifically, Factor 1 constituted of organic nitrogen compounds (Fig. 2A), including oligopeptide and nitrogen-containing alkaloids (such as Ethylmorphine, Levallorphan, and Alangicine), and their abundances were seen consistently declining in the PP group (Table S4). The serum levels of lipids in the PP group (Factor 2), such as Oleoylethanolamide (Fold-change, FC = 158.85), 17beta-Nitro-5alpha-androstane (FC = 59.90), Thiofanox (FC = 54.32), and 17-Methylstearate (FC = 17.33) were observed to be considerably elevated (Fig. 2A, Table S4). In addition, an appreciable reduction in several other differential metabolites, such as organosulfur compound (Factor 3), phenylalanine (Factor 5), and terpenoid (Factor 6) were detected in the PP group as compared to the normal group (Table S4).

### Associations Between Microbial Factors and Metabolic Factors

Subsequently, the associations between microbial and metabolic factors were evaluated to discover the potential key drivers of such modifications. Relatively, organic polycyclic compound expressed vital positive correlations with *Lachnospira*, Enterobacteriaceae, *Dialister*, Peptostreptococcaceae, and Burkholderiaceae (Fig. 2C). Among them, Enterobacteriaceae was positively associated with aferine, phenylalanine, and with LPC as well. The overlapping metabolic pathways, such as amino acid, carbohydrate, aromatic compound metabolic pathway, and others (Glycolysis II, Glyoxylate cycle, and Incomplete reductive TCA cycle), could prove to be the foundation of the strong relationships between microbiota and metabolites (Fig. S1B). Interestingly, even though *Bacteroides* and butyrate-producing bacteria exhibited striking correlations to most metabolic pathways (Fig. S1B), they were not significantly associated with the differential metabolites of PP (Fig. 2C). Furthermore, organic nitrogen compounds and lipids revealed no substantial connection with most of the gut microbiota factors, except for factor *Alistipes* and factor *Dialister* (Fig. 2C), suggesting that they may be directly affected by other factors, such as lifestyles.

### Three PP Subtypes Were Revealed by Differential Metabolites

In comparison to the gut microbiota, more evident modifications were seen in serum metabolism (Fig. S3A, B). The girls in the PP group were classified into three subgroups based on the expression of 182 differential metabolites, all of which demonstrated varied metabolite patterns, implying that PP could be categorized into 3 different subtypes. Specifically, latent metabolic factors, organic nitrogen compounds, and organic polycyclic compounds emerged critically different among the three defined subtypes (Fig. S3C and Table S5). Moreover, some latent microbial factors, such as Peptostrptococcaceae and Lachnospira, also expressed varying abundance patterns among these subtypes. Additionally, clinical laboratory tests, such as for LH, testosterone (TES), Zn, and Ca, presented a similar trend (Fig. S3C). Although the widely utilized standard suggests no difference between the various phenotypes (Table S5), subtype 2 showed a tendency to be distinct from the other two subtypes and appeared to be more analogous to the normal group, which highlights the individual differences among the PP group, especially in the metabolic changes.

### Dietary Pattern Varied Significantly between Groups

Detailed lifestyle information that may potentially affect the PP group was obtained, including dietary patterns, living environment, maternal health, childbirth, and physical condition of the participants and their parents (Table S1). Dietary patterns presented considerable differences among the patients and displayed good discriminative ability for distinguishing the PP group from the normal controls, obtaining an AUC of about 0.87 (Fig. S2D). However, other lifestyle variables failed to express any significant variations between the normal and the PP groups (Table S1) along with a poor discriminative ability (Fig. S2E-H).

Furthermore, EFA facilitated the identification of 9 latent dietary factors derived from the dietary patterns recorded (Fig. 3A, Table S6), which preserved the foremost information of dietary patterns (AUC = 0.86, Fig. 3C). As expected, these latent dietary factors were an amalgamation of the various aspects of the children’s diet, such as healthy foods, junk foods, items containing monosodium glutamate (MSG), and the balance between meat and vegetables. The most critical latent dietary factor, healthy foods, appeared to have a significantly lower intake rate in the PP group (FDR = 0.30e-4, Fig. 3B), which entails preferences for seafood, freshwater products, tubers, vegetables, bean products, fruits, nuts, etc. (Table S6). Several noteworthy correlations were noticed between dietary, microbial, and metabolic factors (Fig. 3D). The intake of healthy foods expressed a highly negative correlation with the abundance of serum lipid (*P* < 0.01). On the other hand, the intake of snacks and drinks were witnessed to be positively linked with the organic nitrogen compounds (*P* < 0.01), whereas negatively correlated with butyrate-producing bacteria (*P* < 0.05). These results were indicative of the unbalanced dietary patterns influencing the PP progression through microbiota and metabolites.

**Fig. 3.**
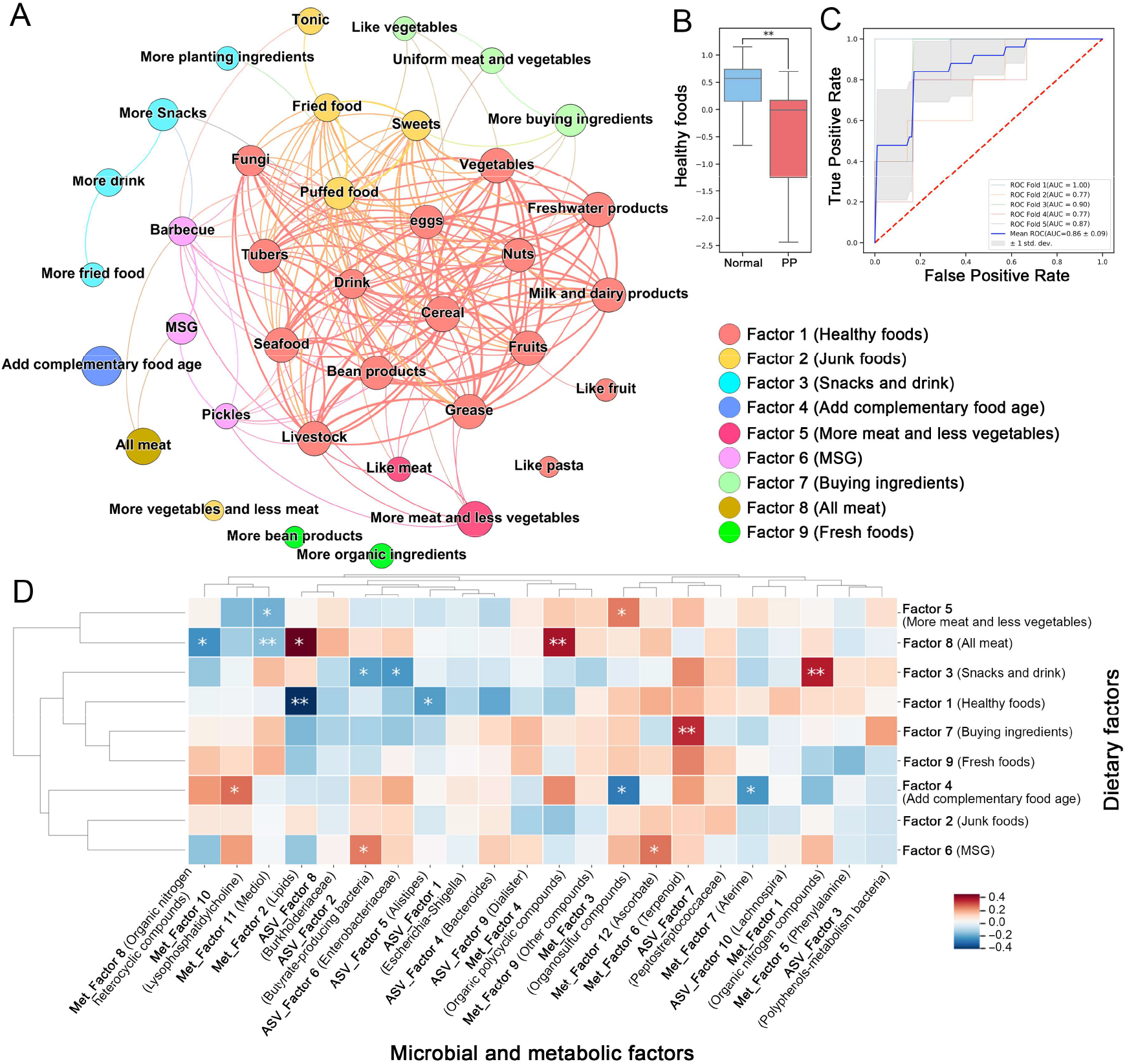
The change of dietary patterns in girls with partial PP. A. Co-occurrence network of dietary patterns through Spearman’s or Kendall’s rank correlation analysis with *P*<0.05. Dietary patterns are colored by their main latent metabolic factor and the strength of correlation is represented by line thickness. B. The difference of healthy food (Factor1) intake between normal and PP groups. C. The ROC curve of the disease discriminating ability with 9 latent dietary factors. D. Heatmap plot of Pearson correlation between latent dietary factors and microbial or metabolic latent factors. (*: *P* < 0.05, **: *P* < 0.01.)

### Unbalanced Dietary Patterns Affecting PP Progression Through Microbiota and Metabolites

Supported by the above results, potential causal relationships among gut microbiota, metabolites, dietary patterns, and the characteristics of the disease were investigated. For this purpose, we introduced the innovative SEM path analysis to construct a credible model (Fisher’s C = 815.85 with *P* = 0.341) in accordance with the correlation results (Fig. 2, 3 and Fig S2I) and our prior knowledge about the subject, to reveal the internal connections among them (Fig. 4). Hormones (estradiol (E2), prolactin (PRL), LH and FSH, and trace elements (Zn, Ca, My, Cu, Fe) were significantly regulated by gut microbiota and serum metabolism. Among the serum metabolism agents, lipids elevated in the most dramatic fashion in the PP group (Table S4), positively affecting the ovarian volume (*P* < 0.05), while producing a negative effect on the breast volume (*P* < 0.05). The reduced intake of healthy foods (*P* < 0.001) and elevated intake of all-meat diet (*P* < 0.001) were known to be the primary factors causing the surge of serum lipids. Similarly, organic nitrogen compounds, the principal metabolic factor, produced a significantly positive effect on the level of serum E2 and further affected the development of ovarian volume. As an important sex hormone, LH was appreciably decreased in the PP group (*P* < 0.01, Table S7) and generated a considerably positive effect on the bone age (*P* < 0.05) and uterine volume (*P* < 0.001) in the PP group (Fig. 4). In serum metabolism, LPC may be the chief influencing factor of LH (*P* < 0.05, Fig. S2I). Furthermore, butyrate-producing bacteria, including genus *Subdoligranulum* and *Faecalibacterium* of family Ruminococcaceae, produced stronger positive effects on the follicle size (*P* < 0.05). The up-regulation of butyrate-producing bacteria may possibly be able to explain the premature development of follicles in the PP group.

**Fig 4.**
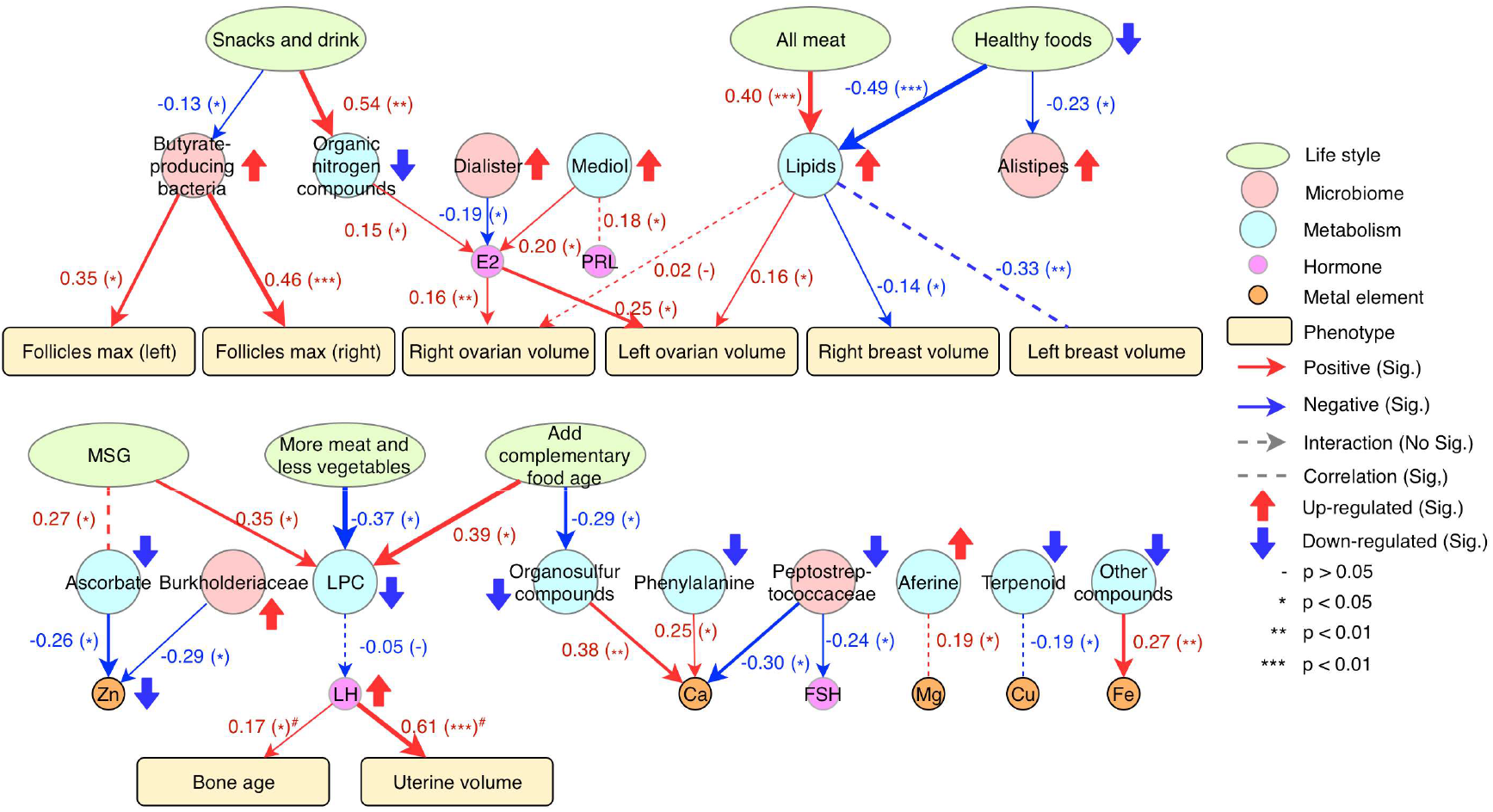
Causal relationships between the latent factors of the dietary, gut microbiota, metabolism, and clinical characteristics. Structural equation model is implemented using piecewise SEM in R. Different types of causal effect are colored by different colors (red: positive, blue: negative) and the strength of effect is represented by line thickness. (-:*P*>0.05, *: *P* < 0.05, **: *P* < 0.01, ***: *P* < 0.001, ^#^: the regression uses glm in R with Poisson distribution).

The SEM analysis revealed that dietary patterns were the most vital of all the catalysts of change in microbiota and metabolism. Simultaneously, the dysbiosis of the gut bacteria taxa and metabolites produced a remarkable effect on the host hormone levels and PP progression (Fig. 4).

## DISCUSSION

Mounting evidence suggests that gut microbiota and metabolism are the major decisive forces of the growth in children (Tamburini et al., 2016a; Yatsunenko et al., 2012). Nevertheless, no definite explanation is available about the effects of gut microbiota and serum metabolism in the pathogenesis of precocious puberty. In this study, we performed a systematic analysis investigating the lifestyle patterns, altered microbiota, metabolome, and their relationship with PP.

It was found that the PP group showed a significant dysbiosis of gut microbiota and serum metabolome, which could be mainly attributed to the unhealthy dietary habits, directly affecting the progression of PP. At the same time, gut microbiota and serum metabolome could prove to be non-invasive as well as reliable diagnostic biomarkers for the early detection of PP, expressing an AUC = 0.93 and 0.97, respectively, and can help evade the time-consuming and painful gonadotropin-releasing hormone (GnRH) stimulation test (Kim et al., 2011a).

Diet shapes our gut microbiota in the early stages of our life (Tamburini et al., 2016b; Zmora et al., 2019). A large number of studies have suggested that unhealthy dietary habits shift the gut microbiota and may very well contribute to the pathogenesis of various metabolic diseases, including overweight, obesity, type 2 diabetes, non-alcoholic liver disease, cardio-metabolic diseases, and malnutrition (Fan and Pedersen, 2020; Tamburini et al., 2016b). An extensively shifted microbial composition was detected in the PP group, implying a dysbiosis of gut bacterial community. β-glucuronidase-producing bacteria, including *Alistipes, Bacteroides, Escherichia*, and *Faecalibacterium* were noticed to be significantly increased in the PP group, possibly triggering the elevated levels of circulating estrogen and increased estrogenic burden, which actuate the onset of PP (Baker et al., 2017; Mcintosh et al.; Sultan et al., 2012). The shift in gut microbiota, especially *Bacteroides* and butyrate-producing bacteria is capable of affecting various metabolic processes (Fan and Pedersen, 2020), such as carbohydrate, fatty acid, and lipid metabolism activity (Fig. S1). Interestingly, butyrate-producing bacteria, which was considered to be a beneficial bacteria for maintaining the gut health in several preceding studies (Cheng et al., 2016; Valles-Colomer et al., 2019), may emerge as a pathogenic agent for PP. Considering the promoting effect of butyrate to the levels of LH (Ruddon et al., 1979) and FSH (Ghosh and Cox; Liang et al., 2020), we propose that overproducing butyrate induced by gut bacteria may produce a detrimental effect on the health condition during puberty, especially for follicular development (Fig. 4). Consistent with the previous study on the gut microbiota of girls suffering from PP conducted by Dong et al., the gut bacteria related to the production of short-chain fatty acids (SCFAs) are known to be present in increased concentrations in girls suffering from PP, which promote the expression of the leptin gene, activate the HPG axis through a high concentration of SCFAs, and trigger the early onset of puberty (Dong et al., 2019).

Moreover, PP gives rise to a more serious metabolic disturbance. Differential metabolites exhibit a strong ability for PP diagnosis (AUC = 0.97) and present three different PP subtypes. A large amount of organic nitrogen compound down-regulation and lipid up-regulation with high fold-change were seen to be the chief characteristics of PP serum metabolome. Decreased intake of healthy foods (*P* < 0.001), the unique differential dietary pattern of PP, illustrated a direct effect on the serum levels of lipids (2-158 fold increase), which was previously verified in high-fat diet mice (Walker et al., 2017). Overnutrition and excessive intake of processed and high-fat food leads to obesity at the beginning of the PP pathogenesis (Latronico et al., 2016; Mugo et al., 2007). In addition, animal studies have also indicated that postnatal overnutrition consistently increases the LH, FSH, leptin, and insulin levels in pubertal females, and postnatal high fat diet after commencing weaning tends to trigger advanced puberty in females [31,32]. The dysbiosis of the serum metabolites indicates an influence on the level of serum metallic elements (Ca, Zn, Cu, Mg, and Pb), which could be potential endocrine disrupters that are capable of modulating estrogenic activity of endogenous hormones (Arjmandi et al., 1993).

Unlike the traditional correlation research, this study adopted the causal inference method based on the SEM analysis, which has been progressively implemented for varied purposes in the microbiome field (Chen et al., 2019; Mamet et al., 2019). This method can effectively derive and comprehend the causal relationships between dietary patterns, gut microbiota, serum metabolome, and PP. Being constrained by prior knowledge and data integrity, the explanation for some results remains unclear, and requires further in-depth research.

In summary, it was found that unhealthy dietary habits could disrupt the homeostasis of gut microbiota and serum metabolism, and consequently trigger the imbalance of hormones, leading to the excessive change of physical development progress and PP genesis. Therefore, the intestinal microbiota may be regarded as a prospective therapeutic target for the prevention and treatment of PP.

### Limitations of Study

The etiology of PP remains complicated; hence it requires further validation by employing larger samples and effectively designing disease prediction models. In addition, although this study included the validation cohorts, the multi-center studies and big sizes of the validation cohorts will be needed to further validate the biomarkers found in this study.

## Materials and methods

### Study participants

133 girl participants with partial PP and 67 normal girls were recruited in the discovery cohort at Qilu Children’s Hospital, Shandong University (Table 1) for this study. 66 normal and 17 girls suffering from PP (age < 8) were included in the independent validation cohort to externally verify our findings. The PP group was diagnosed in accordance with the criteria defined by Lawson Wilkins Pediatric Endocrine Society. Exclusion criteria includes (1) other organic etiologies with presence of isointense tumor on magnetic resonance imaging (MRI); (2) usage of antibiotics, probiotics, or prebiotics within 3 months before enrolling; (3) associated endocrine, gastrointestinal, metabolic disease (including obesity and diabetes, among others), mental disease, or hepatobiliary disease. Recruited age-matched normal controls were girls below 8 years of age without PP. The study protocol was maintained in accordance with the Declaration of Helsinki and was approved by the Ethics Committee of Qilu Children’s Hospital (ETYY-2016-202). Written informed consents and questionnaires were obtained from the children’s parents.

### Sample collection

Stool and blood samples were collected from each participant and stored at -80 °C before analysis. 200 mg stool was preserved in sterile 2 ml tubes containing pure ethanol, aliquoted (Tinygene Biological Company, China) and stored at -80 °C for 16S rRNA sequencing. Blood samples were thawed at 4 °C, 3000 rpm, and centrifuged at 4 °C for 10 min. Serum aliquots were immediately frozen at -80 °C for further untargeted metabolomics analysis. The study was approved by local ethics committees (Qilu Children’s Hospital of Shandong University, IRB Number EYY-2016-202) and informed consent was obtained from all the participating subjects.

### DNA extraction and illumina sequencing

Total DNA extraction from fecal samples (250 mg, wet weight) was performed using a Fast DNA SPIN Kit for feces (MP Biomedicals, Santa Ana, CA, USA), as per the manufacturer’s instructions. The V1-V2 hypervariable region was amplified with the universal primer pair F27 (5’ -AGAGTTTGATCMTGGCTCAG-3’) and R355 (5’-GCTGCCTCCCGTAGGAGT -3’). Sequencing was conducted on Illumina HiSeq 2500 System (Illumina Inc., San Diego, CA, USA) using the 2 × 250 paired-end mode following the standard Illumina platform protocols. All sequencing data is available at NODE (http://www.biosino.org/node) with the accession number OEP000731.

### 16S rRNA gene sequencing data analysis

16S rRNA sequencing data was analyzed using Quantitative Insights Into Microbial Ecology (QIIME2 V.2019.07). In brief, raw sequence data was demultiplexed and DADA2 (SP) was employed to denoise sequencing reads for quality control and the identification of amplicon sequence variants (ASVs) via q2-dada2 plugin. Taxonomy classification was carried out by utilizing classify-sklearn based on a Naiva Bayes classifier against the Silva-132-99 reference sequences. Respective sequences of each ASV were aligned with Multiple Alignment using Fast Fourier Transform (MAFFT) (Katoh et al., 2002) (via q2-alignment) and the phylogenetic tree was constructed with Fast-Tree (Price et al., 2010) (via q2-phylogeny). Chao1, an index of richness estimator, was calculated to assess the community alpha diversity. Principal coordinate analysis (PCoA) was performed based on the Bray-Curtis distance; and PERMANOVA test was conducted to evaluate the significant differences present among the microbial communities, with 9999 permutations.

Additionally, the functions of gut microbiota were inferred based on 16S rRNA sequencing data using Phylogenetic Investigation of Communities by Reconstruction of Unobserved States (PICRUSt2), as previously described (Douglas et al., 2020).

### Untargeted metabolomics and analysis

100 μL of serum sample was transferred to an EP tube. After the addition of 300 μL of methanol (containing internal standard 1 μg/mL), the samples were vortexed for 30 s, followed by sonication for 10 min in ice-water bath, and incubation for 1 h at -20 °C to precipitate the proteins. The sample was then centrifuged at 12000 rpm for 15 min at 4 °C. The resulting supernatants were then transferred to LC-MS vials and stored at -80 °C until the UHPLC-QE Orbitrap/MS analysis. The quality control sample was prepared by mixing an equal aliquot of the supernatants from all the samples collected.

LC-MS/MS analyses were performed using an UHPLC system (1290, Agilent Technologies) with a UPLC HSS T3 column (2.1 mm × 100 mm, 1.8 μm) coupled to Q Exactive (Orbitrap MS, Thermo). The mobile phase A was 0.1% formic acid in water for positive, and 5 mmol/L ammonium acetate in water for negative, and the mobile phase B was acetonitrile. The elution gradient was set as follows: 0 min, 1% B; 1 min, 1% B; 8 min, 99% B; 10 min, 99% B; 10.1 min, 1% B; 12 min, 1% B. The flow rate was 0.5 mL/min. The injection volume was 2 μL. The QE mass spectrometer was utilized due to its ability to acquire MS/MS spectra on an information-dependent basis (IDA) during an LC/MS experiment. In this mode, the acquisition software (Xcalibur 4.0.27, Thermo) continuously examines the full scan survey MS data as it collects and triggers the acquisition of MS/MS spectra depending on preselected criteria. ESI source conditions were set as follows: Sheath gas flow rate as 45 Arb, Aux gas flow rate as 15Arb, Capillary temperature at 400 °C, Full ms resolution as 70000, MS/MS resolution as 17500, Collision energy as 20/40/60 eV in NCE model, Spray Voltage as 4.0 kV (positive) or -3.6 kV (negative), respectively.

Raw data was converted to mzXML format using ProteoWizard and processed using MAPS software (version 1.0). Preprocessed results were employed to generate a data matrix which consisted of the retention time (RT), mass-to-charge ratio (m/z) values, and peak intensity. In-house MS2 database was applied for metabolites identification.

### Questionnaire survey

Questionnaires were distributed to the participants and their parents in presence of trained doctors, providing professional guidance throughout the whole process. From the questionnaire survey, 117 variables were collected, involving dietary pattern, living environment, maternal health, childbirth, and personal physical condition of the recruited girls (PP and normal group), as well as the physical condition of their parents.

The dietary pattern section in our questionnaire was based on the most frequently consumed foods that were clinically considered to be closely related to PP, including vegetables, fruits, seafood, meat, cereal, tubers, eggs, milk, bean products, nuts, fungi, greasy food, beverages, fried food, sweets, barbecued food, puffed food, pickles, gourmet powder, and tonic, on a 4-level intake frequency scale. The dietary section also entailed personal food preferences, including preferences for fruit, vegetables, meat, pasta, bean products, fried food, snacks, beverage, meat, etc.

The variables regarding living environment included presence of pets, use of mineral water, existence of bowel dysfunction of close contacts, presence of plastic foam products, pesticides, fertilizers, insecticide, etc. The questionnaire also separately covered the physical condition of participants’ parents, including stomachaches, abdomen distension, diarrhea, gastric acid regurgitation, constipation, hypertension, hyperglycemia, anemia, rheumatism, urticaria, immunodeficiency, and irregular menstruation and menarche age (only for mothers). The data about maternal conditions during pregnancy and delivery was derived, including reproductive age, medication during pregnancy, secretory disorders during pregnancy, folic acid supplement, dietary patterns, alcohol consumption, abnormal fetal movement, oxytocin, dystocia, or fetal hypoxia, cesarean delivery, or spontaneous delivery, etc. The physical condition of the recruited girls was investigated thoroughly, including sleep disorder, poor weight gain, jaundice, eczema, diarrhea, oral malodor, sediment in urine, smelly urine, perianal red, dental caries, etc.

### Clinical laboratory tests

Clinical parameters were determined at the clinical lab of Qilu Children’s Hospital. The trace element (Cu, Zn, Ca, Mg, Fe, and Pb) levels from serum samples were measured using the flame atomic absorption method (BH5100, Bohui, China). The thyroid function test was conducted by analyzing the serum levels of follicle stimulating hormone (FSH), luteinizing hormone (LH), and thyroid stimulating hormone (TSH), employing the chemi-luminescence immunoassay methods (Abbott, Architect I2000, US). GnRH stimulation test was conducted for the luteinizing hormone (LH) and follicle-stimulating hormone (FSH) utilizing the chemi-luminescence immunoassay methods (Abbott, Architect I2000, US). Cortisol (COR), adrenocortieotropic hormone (ACTH), alpha fetoprotein (AFP), and carcinoembryonic antigens (CEA) were measured by chemi-luminescence immunoassay methods (LIAISON, type 2210, Germany). The plasma levels of the insulin and insulin-like growth factors were quantified by adopting immunoluminescence method (Siemens immulite 2000, USA). The sizes of uterus, breasts, and ovaries, as well as the number of ovarian follicles were determined by employing B-ultrasonic examination (EPIQ5, L12-5, Philips, Holland). The hand-wrist radiographs were used for bone-age assessment through nuclear magnetic resonance (MRI) examination (Digital Dianost3, Philips, Holland).

### Co-occurrence analysis

Co-occurrence analysis was applied for microbial, metabolic, or dietary network by using correlations (Spearman, Spearman or Kendall). Correlations with *P* ≤ 0.05 (permutation test with 1000 permutations) were included in the co-occurrence networks. Network visualization was conducted using Gephi software (M et al., 2009).

### Disease diagnosis model

Classification model of different samples was constructed using Random Forest classifier in Scikit-learn package of Python (3.6.0)(F et al., 2011). The AUC of 5-fold cross-validation was utilized to measure the discriminative ability of the differential biomarkers.

### Exploratory factor analysis

For differential microbiota, metabolites, and dietary patterns, exploratory factor analysis (EFA) was employed to identify the latent factors with FactorAnalyzer in Python (C, 2016). The number of factors, solutions (minimum residual, maximum likelihood or principal factor), and rotations (varimax or promax) of EFA were determined by minimizing information loss after dimension reduction, which was evaluated by the discriminative ability of the classification of normal and PP samples. Latent factors with high loading were explained and labeled based on the observed variables and former knowledge.

### Structural equation model

Based on the latent factors from EFA, path analysis of the SEM (D et al., 2019) was employed to discover the causal relationships between lifestyle, gut microbiota, metabolism, and clinical characteristics of disease. Considering that the samples cannot match perfectly, piecewise SEM (Shipley, 2013) was used for confirmatory path analysis in our study, in which each set of relationships was determined independently (or locally). The p-value of Fisher’s C statistic was adopted to prove the overall rationality of the model and to facilitate the model comparison and selection.

### Statistical analysis

All statistical analyses were conducted using Python (3.6.0). Statistical significance was determined by two-sided Wilcoxon rank-sum test, Permutation test or one-sided Fisher’s exact test, and Benjamini-Hochberg test was applied to control the false positive rate (FDR) under multiple comparisons. Differences were considered statistically significant when FDR < 0.05.

## Author contributions

Zhongtao Gai, Ruixin Zhu, Lei Zhang, Guoping Zhao and Guoqing Zhang conceived and designed the project. Each author has contributed significantly to the work submitted. Hongying Li, Xiangrong Liang, Qian Wang, Meiling Huo, Ying Wang, Lu Zhao, Yongsheng Ge, and Changying Zhao performed the clinical diagnosis, designed and collected the clinical setting, underwent the ethical evaluation process, discussed informed consent and questionnaire data sheets with patients. Ying Wang, Lu Zhao, Yongsheng Ge, and Changying Zhao collected clinical samples and patient information, and consolidated and organized the whole data. Xin Lv completed the clinical examination work and provided the relevant data. Dingfeng Wu and Na Jiao analyzed the data. Ying Wang and Dingfeng Wu drafted the manuscript. Na Jiao, Wenxing Gao, Sheng Gao, Liwen Tao, Ruifang Cao, Yunchao Ling, Lingna Zhao, Haokui Zhou, Guoqing Zhang, Guoping Zhao, Lei Zhang, Ruixin Zhu and Zhongtao Gai revised the manuscript. All the authors read and approved the final manuscript.

## Supplementary materials

**Fig. S1.**
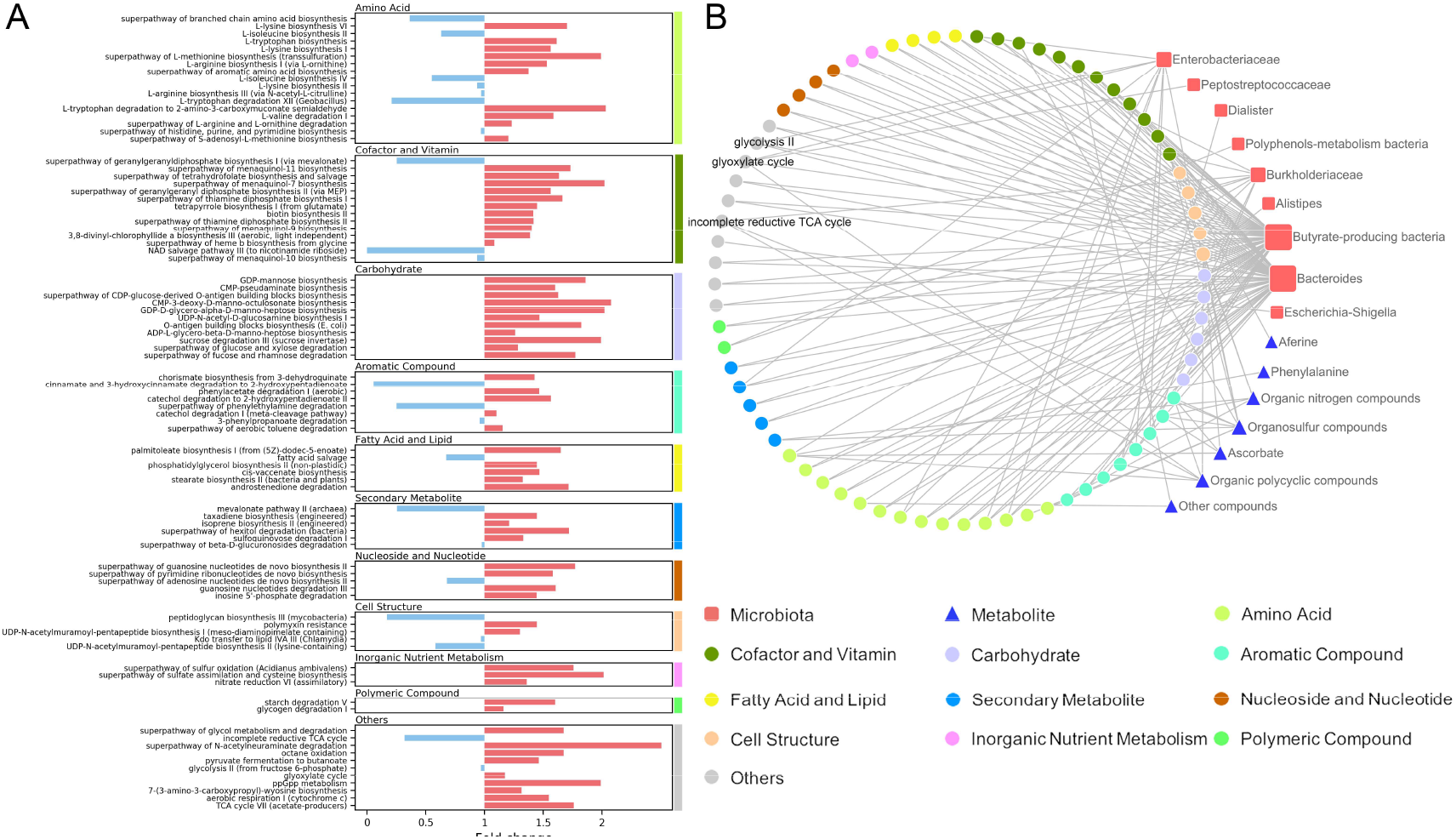
The dysbiosis of metabolic pathways in gut microbiota. A. The differential abundance changes of gut microbial metabolic pathways. B. The significant correlation links between gut microbiota, serum metabolome, and pathways.

**Fig. S2.**
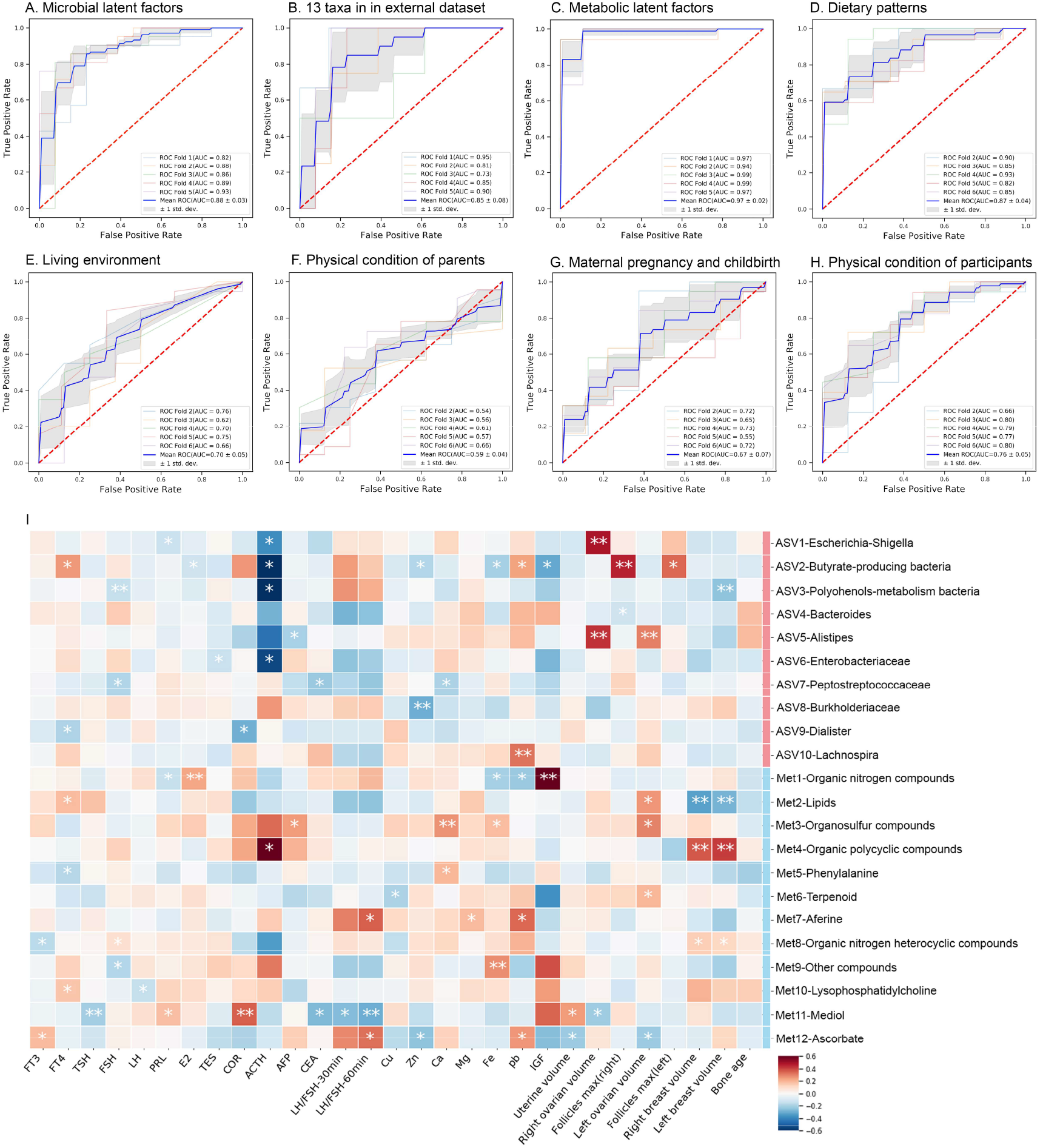
The ROC curve of the disease discriminating ability and Correlation between latent microbial factors, latent metabolic factors, hormones, and phenotypes. The ROC curve of the disease discriminating ability with 10 latent microbial factors (A), 13 taxa in external dataset (B), 12 metabolic latent factors (C), dietary patterns (D), living environment (E), physical condition of parents (F), maternal health and childbirth (G) and physical condition of participants (H). I. Correlation between latent microbial factors, latent metabolic factors, hormones, and phenotypes. All relationships were calculated by Pearson correlation in Python with 1000 permutation test (*:*P* < 0.05, **: *P* < 0.01.)

**Fig. S3.**
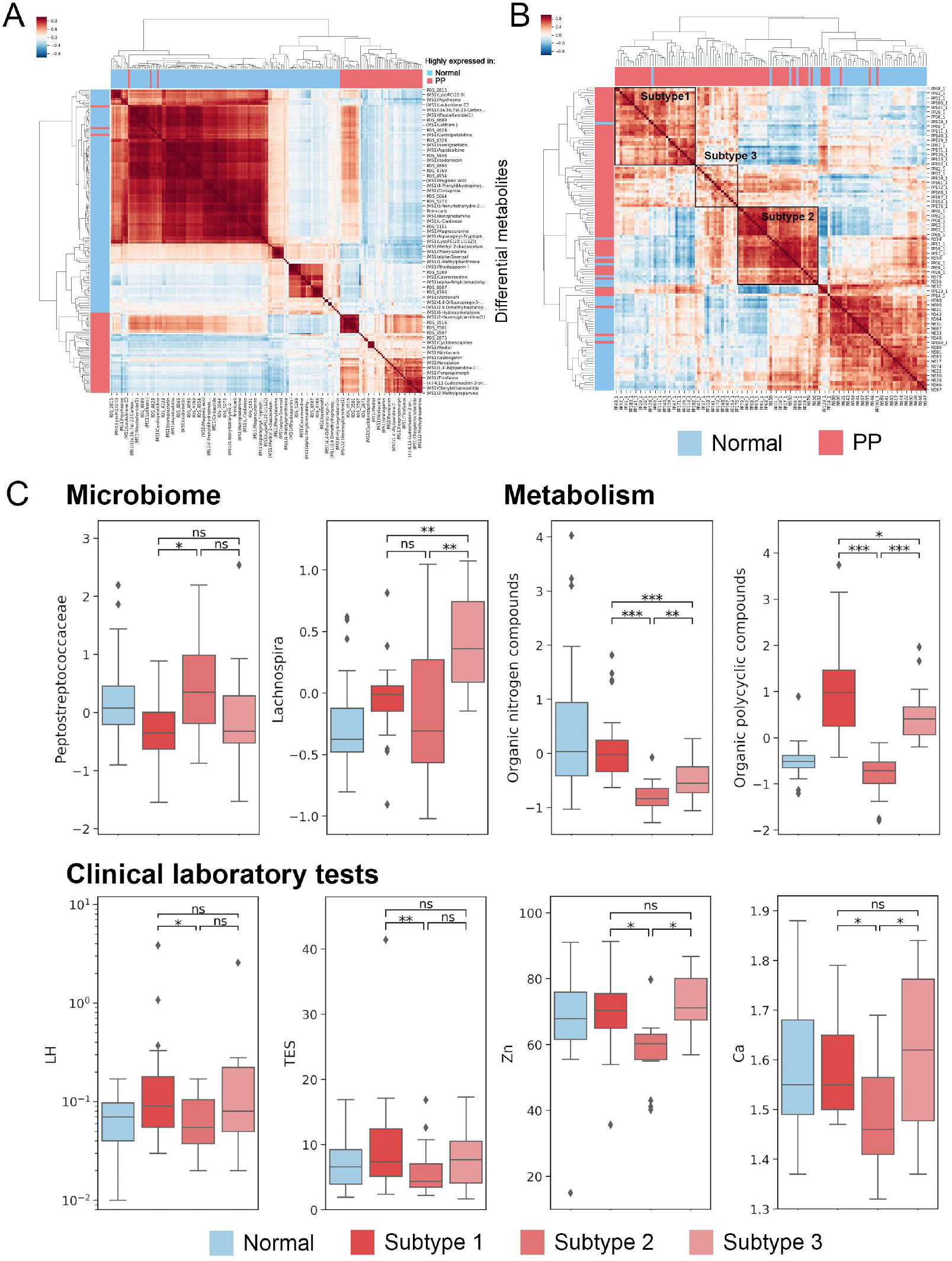
Three different subtypes in PP group. A. Pearson correlation between differential metabolites. B. Pearson correlation of samples with differential serum metabolites. Three subtypes were defined through hierarchical clustering algorithm with Euclidean distance. C. The difference between subtypes in gut microbiota, serum metabolome, and clinical laboratory tests. (ns: *P*>0.05, *: *P* < 0.05, **: *P* < 0.01, ***: *P* < 0.001.).

**Table S1. Differential life styles between normal and PP group**.

**Table S2. Differential gut microbiota between normal and PP group and latent factors**.

**Table S3. 13 taxa presented same differential abundance change in external dataset**.

**Table S4. Differential serum metabolites between normal and PP group and latent factors**.

**Table S5. Gut microbiota, serum metabolome, clinical laboratory tests, and phenotype difference between PP subtypes**.

**Table S6. Latent factors of life styles**.

**Table S7. Differential clinical laboratory variables between normal and PP group**.

## REFERENCES

Arjmandi, B.H., Salih, M.A., Herbert, D.C., Sims, S.H., and Kalu, D.N. (1993). Evidence for estrogen receptor-linked calcium transport in the intestine. Bone and mineral 21, 63–74.

Baker, J.M., Al-Nakkash, L., and Herbst-Kralovetz, M.M. (2017). Estrogen–gut microbiome axis: physiological and clinical implications. Maturitas 103, 45–53.

Brito, V., Batista, M., Borges, M., Latronico, A., Kohek, M., Thirone, A., Jorge, B., Arnhold, I., and Mendonca, B. (1999). Diagnostic value of fluorometric assays in the evaluation of precocious puberty. The Journal of Clinical Endocrinology & Metabolism 84, 3539–3544.

C, H.M. (2016). A review of exploratory factor analysis decisions and overview of current practices: What we are doing and how can we improve? International Journal of Human-Computer Interaction 32, 51–62.

Chen, C., Chen, Y., Zhang, Y., Sun, W., Jiang, Y., Song, Y., Zhu, Q., Mei, H., Wang, X., and Liu, S. (2018). Association between dietary patterns and precocious puberty in children: a population-based study. International journal of endocrinology 2018.

Chen, L., Jiang, Y., Liang, C., Luo, Y., Xu, Q., Han, C., Zhao, Q., and Sun, B.J.M. (2019). Competitive interaction with keystone taxa induced negative priming under biochar amendments. 7, 1–18.

Cheng, J., Ringel-Kulka, T., Heikamp-de Jong, I., Ringel, Y., Carroll, I., de Vos, W.M., Salojarvi, J., and Satokari, R. (2016). Discordant temporal development of bacterial phyla and the emergence of core in the fecal microbiota of young children. ISME J 10, 1002–1014.

Cussotto, S., Sandhu, K.V., Dinan, T.G., and Cryan, J.F. (2018). The neuroendocrinology of the microbiota-gut-brain axis: a behavioural perspective. Frontiers in neuroendocrinology 51, 80–101.

D, M.S., E, R., M, B., G, L.E., L, H.B., K, S., and D, S.S. (2019). Structural equation modeling of a winnowed soil microbiome identifies how invasive plants re-structure microbial networks. The ISME journal 13, 1988–1996.

Dong, G., Zhang, J., Yang, Z., Feng, X., Li, J., Li, D., Huang, M., Li, Y., Qiu, M., and Lu, X. (2019). The association of gut microbiota with idiopathic central precocious puberty in girls. Frontiers in Endocrinology 10.

Douglas, G.M., Maffei, V.J., Zaneveld, J.R., Yurgel, S.N., Brown, J.R., Taylor, C.M., Huttenhower, C., and Langille, M.G.J.N.B. (2020). PICRUSt2 for prediction of metagenome functions. 1–5.

Du, G., Hu, J., Huang, Z., Yu, M., Lu, C., Wang, X., and Wu, D. (2019). Neonatal and juvenile exposure to perfluorooctanoate (PFOA) and perfluorooctane sulfonate (PFOS): Advance puberty onset and kisspeptin system disturbance in female rats. Ecotoxicology and environmental safety 167, 412–421.

F, P., G, V., A, G., V, M., B, T., O, G., M, B., P, P., R, W., V, D., et al. (2011). Scikit-learn: Machine learning in Python. the Journal of machine Learning research 12, 2825–2830.

Fan, Y., and Pedersen, O. (2020). Gut microbiota in human metabolic health and disease. Nature Reviews Microbiology, 1–17.

Frankenfeld, C.L., Atkinson, C., Wähälä, K., and Lampe, J.W. (2014). Obesity prevalence in relation to gut microbial environments capable of producing equol or O-desmethylangolensin from the isoflavone daidzein. 68, 526–530.

Ghosh, N.K., and Cox, R.P. Induction of human follicle-stimulating hormone in HeLa cells by sodium butyrate. Nature 267, 435–437.

Katoh, K., Misawa, K., Kuma, K.i., and Miyata, T. (2002). MAFFT: a novel method for rapid multiple sequence alignment based on fast Fourier transform. Nucleic acids research 30, 3059–3066.

Kim, H.K., Kee, S.J., Seo, J.Y., Yang, E.M., Chae, H.J., and Kim, C.J. (2011a). Gonadotropin-releasing hormone stimulation test for precocious puberty. The Korean journal of laboratory medicine 31, 244–249.

Kim, J., Kim, S., Huh, K., Kim, Y., Joung, H., and Park, M. (2011b). High serum isoflavone concentrations are associated with the risk of precocious puberty in Korean girls. Clinical endocrinology 75, 831–835.

Kim, S.H., Huh, K., Won, S., Lee, K.-W., and Park, M.-J. (2015). A significant increase in the incidence of central precocious puberty among Korean girls from 2004 to 2010. PLoS One 10.

Kong, L.C., Holmes, B.A., Cotillard, A., Habi-Rachedi, F., Brazeilles, R., Gougis, S., Gausserès, N., Cani, P.D., Fellahi, S., and Bastard, J.-P. (2014). Dietary patterns differently associate with inflammation and gut microbiota in overweight and obese subjects. PloS one 9.

Latronico, A.C., Brito, V.N., and Carel, J.-C. (2016). Causes, diagnosis, and treatment of central precocious puberty.The Lancet Diabetes & Endocrinology 4, 265–274.

Liang, Y., Ming, Q., Liang, J., Zhang, Y., Zhang, H., and Shen, T. (2020). Gut microbiota dysbiosis in Polycystic ovary syndrome (PCOS): association with obesity - A preliminary report. Canadian Journal of Physiology and Pharmacology.

M, B., S, H., and M, J. (2009). Gephi: an open source software for exploring and manipulating networks. Third international AAAI conference on weblogs and social media.

Mamet, S.D., Redlick, E., Brabant, M., Lamb, E.G., Helgason, B.L., Stanley, K., and Siciliano, S.D.J.T.I.j. (2019). Structural equation modeling of a winnowed soil microbiome identifies how invasive plants re-structure microbial networks. 13, 1988–1996.

Mcintosh, F.M., Maison, N., Holtrop, G., Young, P., Stevens, V.J., Ince, J., Johnstone, A.M., Lobley, G.E., Flint, H.J., and Louis, P. Phylogenetic distribution of genes encoding β-glucuronidase activity in human colonic bacteria and the impact of diet on faecal glycosidase activities. Environmental Microbiology 14, 0–0.

Merzenich, H., Boeing, H., and Wahrendorf, J. (1993). Dietary fat and sports activity as determinants for age at menarche. American journal of epidemiology 138, 217–224.

Mugo, M.N., Bachrach, B.E., and Jacobson, J.D. (2007). Partial and Gonadotropin-Dependent Precocious Puberty.

Muir, A. (2006). Precocious puberty. Pediatrics in Review 27, 373.

Price, M.N., Dehal, P.S., and Arkin, A.P. (2010). FastTree 2–approximately maximum-likelihood trees for large alignments. PloS one 5.

Qi, Y., Li, P., Zhang, Y., Cui, L., Guo, Z., Xie, G., Su, M., Li, X., Zheng, X., and Qiu, Y. (2012). Urinary metabolite markers of precocious puberty. Molecular & Cellular Proteomics 11.

Rogers, I.S., Northstone, K., Dunger, D.B., Cooper, A.R., Ness, A.R., and Emmett, P.M. (2010). Diet throughout childhood and age at menarche in a contemporary cohort of British girls. Public health nutrition 13, 2052–2063.

Root, A.W. (2000). Precocious puberty. Pediatrics in review 21, 10–19.

Ruddon, R.W., Anderson, C., Meade, K.S., Aldenderfer, P.H., and Neuwald, P.D. (1979). Content of gonadotropins in cultured human malignant cells and effects of sodium butyrate treatment on gonadotropin secretion by HeLa cells. Cancer Research 39, 3885–3892.

Sheflin, A.M., Melby, C.L., Carbonero, F., and Weir, T.L. (2017). Linking dietary patterns with gut microbial composition and function. Gut Microbes 8, 113–129.

Shipley, B. (2013). The AIC model selection method applied to path analytic models compared using a d-separation test. Ecology 94, 560–564.

Soliman, A., De Sanctis, V., and Elalaily, R. (2014). Nutrition and pubertal development. Indian journal of endocrinology and metabolism 18, S39.

SP, D. High-resolution sample inference from Illumina amplicon data. (Nat).

Sultan, C., Gaspari, L., Kalfa, N., and Paris, F. (2012). Clinical Expression of Precocious Puberty in Girls. Endocr Dev 22, 84–100.

Tamburini, S., Shen, N., Wu, H.C., and Clemente, J.C. (2016a). The microbiome in early life: implications for health outcomes. Nat Med 22, 713–722.

Tamburini, S., Shen, N., Wu, H.C., and Clemente, J.C.J.N.m. (2016b). The microbiome in early life: implications for health outcomes. 22, 713–722.

Valles-Colomer, M., Falony, G., Darzi, Y., Tigchelaar, E.F., Wang, J., Tito, R.Y., Schiweck, C., Kurilshikov, A., Joossens, M., Wijmenga, C., et al. (2019). The neuroactive potential of the human gut microbiota in quality of life and depression. Nat Microbiol 4, 623–632.

Walker, A., Pfitzner, B., Harir, M., Schaubeck, M., Calasan, J., Heinzmann, S.S., Turaev, D., Rattei, T., Endesfelder, D., and zu Castell, W. (2017). Sulfonolipids as novel metabolite markers of Alistipes and Odoribacter affected by high-fat diets. Scientific reports 7, 1–10.

Yatsunenko, T., Rey, F.E., Manary, M.J., Trehan, I., Dominguez-Bello, M.G., Contreras, M., Magris, M., Hidalgo, G., Baldassano, R.N., Anokhin, A.P., et al. (2012). Human gut microbiome viewed across age and geography. Nature 486, 222–227.

Zmora, N., Suez, J., Elinav, E.J.N.r.G., and hepatology (2019). You are what you eat: diet, health and the gut microbiota. 16, 35–56.

